# Impact of Spruce Plantation on Plant Diversity

**DOI:** 10.1101/2024.02.22.581586

**Authors:** Vanessa Manuzi, Simone Balestra, Pietro Gatti, Gianalberto Losapio

**Affiliations:** Department of Biosciences, University of Milan, Milan, Italy; Institute of Earth Surface Dynamics, Faculty of Geoscience and Environment, University of Lausanne, Lausanne, Switzerland

## Abstract

As the effects of climate change are becoming more evident, different countries around the world are adopting new policies to intervene on the regulation of greenhouse gasses emission. Recent frameworks acknowledge the potential contribution that forest ecosystems can give to carbon sequestration. These are indicating reforestation programmes as effective climate change mitigation options. Yet, there are possibilities that reforestation may have counteractive effects on biodiversity. However the long term consequences of reforestation for biodiversity are poorly understood.

Reforestation policies have already been widely implemented around the world. For instance, in northern Italy and central Europe plantations of spruce trees (*Picea abies*) have been highly promoted during the last century. The objective of our research is to address the long term consequences of reforestation by answering the following questions. What is the spruce plantation’s impact on plant diversity? Does the spruce plantation impact environmental factors like luminosity and ground surface temperature and do these environmental factors affect plant diversity? We hypothesize that the spruce plantation causes a reduction of plant diversity. Indeed, we expect that the spruce plantation affects different environmental factors that have an important role in determining plant composition and abundance.

To answer our research questions, we have conducted a study in two different sites of the Como Prealps. The potential vegetation of the selected area is represented by mixed forests of deciduous trees dominated by beech trees (*Fagus sylvatica*). Historically, the land has also been used for grazing and mowing. However, some stands of the potential vegetation are here replaced by spruce plantations, the presence of which is linked to national forestry policies of the twentieth century. For our research we have conducted a total of 100 vegetation surveys to collect data on plant diversity and environmental factors, namely luminosity and ground surface temperature. We then compared plant diversity among land-use treatments (i.e., habitat types): the spruce plantation, the natural mixed forest and the semi-natural grassland-pasture. For our analysis we have used linear regression models to test the impact of the different habitat types on plant diversity. We have also measured covariance and correlation to analyse the relationship between the environmental factors and plant diversity.

The analysis on plant diversity reveals the long-lasting impact of spruce monoculture plantation on plant diversity. The number of plant species decreases by 57percent from the grassland-pasture to the spruce plantation and by 41percent compared to the natural mixed forest. Likewise, the diversity of plant functional forms decreases in the spruce plantation as compared to mixed forests and grassland-pasture. At last, although luminosity and ground surface temperature do not vary from the mixed forest to the spruce plantation, we have measured a positive relationship between the number of plant species and the two environmental parameters.

Our research provides novel evidence that the spruce plantation negatively impacts plant diversity still one hundred years after. As biodiversity loss and climate change are two interwoven processes, they must not be treated separately. For what concerns future reforestation programmes, we recommend that they include biodiversity-friendly measures and address win-win solutions, for their effectiveness in climate change mitigation would otherwise be compromised.

## 2 Introduction

### 2.1 Mitigating Climate Change with Reforestation

As the effects of climate change are becoming every year more evident [1, 2], concerns on carbon emissions are increasing between governments around the world. The urge to limit greenhouse gasses emission and to stop global warming have been more than once debated by the United Nation Framework Convention on Climate Change (UN-FCCC). In 1997 the Kyoto Protocol was adopted by 192 Parties and later, in 2015, a total of 196 Parties have agreed to sign the Paris Agreement [3, 4]. The UNFCCC acknowledges the twofold influence that forests have as sink for vast amounts of carbon dioxide and, when destroyed or damaged, as source of greenhouse gas emissions. As a consequence, the “REDD+” framework has been established as part of the Paris Agreement. “REDD” stands for “Reducing emissions from deforestation and forest degradation in developing countries”. This framework aims to protect forests and encourage a sustainable management of forest-related activities. [5].

Plants’ ability of absorbing carbon from the atmosphere and creating biomass carbon stocks is well known[6, 7]. As a fact, existing forestland and harvested wood products already contribute in mitigating climate change. Nevertheless, their contribution remains limited as other human activities constantly emit more and more green house gasses. What can help accelerate live-tree sequestration of carbon stocks in forests and the accumulation of carbon in soils is increasing forest area [8, 9]. Not surprisingly, the Clean Development Mechanism (CDM) introduced with the Kyoto Protocol focuses on land management and prioritizes afforestation and reforestation projects[10]. Afforestation involves planting trees where previously there have been none, or where forests have been missing for a long time (50 years according to UNFCCC)[11], while reforestation applies to more recently deforested land[11]. The potential sites that have been identified for the projects’ implementation are pasture lands, mountainous areas, landfills and mined lands[10]. Yet, afforestation and reforestation may have counteractive effects on biodiversity, but the long term consequences of afforestation and reforestation remain poorly understood.

Reforestation and afforestation are considered relatively cost-effective climate change mitigation options[12], however it is likely that planting trees will not be an immediate and permanent solution. Especially on a large scale and in the long term, reforestation and afforestation imply the removal of other habitats from the ecosystems. The removal of habitats may exclude certain groups of species and lead to homogeneous ecosystems, which in turn impact biodiversity and the ecosystems’ functionality[13]. Therefore, before carrying out these reforestation/afforestation projects we need to investigate what their impact on biodiversity is. Filling this knowledge gap will allow the adoption of precautions and alternative measures that can provide long lasting solutions against climate change and, at the same time, limit biodiversity loss while ensuring maintenance of adequate levels of biodiversity and healthy ecosystems.

### 2.2 Types of Reforestation

Today’s Clean Development Mechanism projects are not the first reforestation programmes to have ever taken place in the world. Many of the landscapes that we observe today are the result of centuries and millennia of land management and resource exploitation operated by mankind and societies. In the mid-nineteenth century, most developing countries suffered from great deforestation. The causes behind forest clearing were related to the demands of the growing population, extensive grazing, timber supply for the industry, mining or metallurgical activities, as well as successive wars and fires[14, 15]. Awareness on the consequences of deforestation on hydro-geological risk has lead to reforestation initiatives. As a result, vast areas in different countries are today covered with monocultures, which are also, in some cases, dominated by exotic species[16].

For instance, from the end of the nineteenth century, Spain has undertaken a large-scale afforestation and reforestation process[14]. During this process, *Pinus* species have been widely used because they are characterized by rapid growth and high adaptability to different types of soil. Almost simultaneously, the Australian plant genus *Eucalyptus* has been introduced in Spain and eucalyptus plantations have been widely spread around the country to compensate the mine companies demand of fuel wood[17]. Today, pine forests occupy 32.6 percent of the total forested area in Spain and are the most extended forest type in the country[18]. Eucalyptus plantations represent 3.74 percent of Spain’s forests[17]. As well as in Spain, coniferous monocultures have been extensively planted in other countries of central and northern Europe. The most important tree species of the central European plantations is the Norway spruce (*Picea abies*). Favoured by its ability to grow on many soil types with profitable biomass accumulation and by its wood’s suitability for a variety of purposes, the Norway spruce has been highly used in large-scale afforestation projects practiced from the nineteenth century onward[19]. Today the Norway spruce is locally naturalised in many areas of Europe outside it’s native range and its extreme diffusion has altered the European landscape and its natural forests’ tree composition[15, 20]. Large-scale plantations and introduction of exotic species have occurred outside of Europe too. A well-Known case is Chile’s forest policy established by Pinochet in 1974. Under the pretext of reducing hydro-geological instability, large plantations of *Pinus radiata* and *Eucalyptus spp.* have been used to cover vast areas of the South-central Chilean region, replacing other natural habitats[21].

Nowadays, awareness on the long term consequences of these types of plantations has risen. Not only the expansion of artificial forests - i.e., planted by man[22] - has created less heterogeneous ecosystems, but we presume that their presence also influences other habitat parameters like temperature, humidity and luminosity. In addition to this, we know that coniferous plantation leads to an increase in the acidity of soils[23]. Provided that monocultures cause the alteration of all these different parameters it is likely that they will also affect biodiversity.

Biodiversity plays an essential role in the ecosystem’s regulating and supporting services [24], which include maintaining soil, water and air quality[25], and on the ecosystem’s productivity [13]. As a consequence, biodiversity loss threatens the ecosystem’s functionality, productivity and structure. If an ecosystem loses its complexity beyond a certain limit, it also loses its ability to recover from a disturbance and will collapse[13]. For these reasons, before carrying out new plantation projects to contrast climate change, we must first investigate the impact that already existing plantations have on biodiversity. It is from this urge to find biodiversity-based climate mitigation strategies that our study in the spruce plantations of the Como Prealps has been conceived.

### 2.3 Spruce Monocultures in the Lombardian Prealps

The main objective of this study is to measure how the coniferous monoculture affects biodiversity. For this purpose, we are comparing spruce plantation stands with neighboring mixed forest stands and grassland-pasture stands. In Lombardy the coniferous plantations related to recent reforestation programmes cover a total of 6931 hectares of land[26]. Various spruce plantations are located in the Como Prealps which is where we have selected the sites for our research.

The forests that occupy the Italian territories today result from a long history of human landscape-transforming activities, indeed the woods have been widely exploited and altered by men since ancient times[27]. However, the spruce plantations that occupy the Lombardian Prealps today are a trace of more recent events that have been highly influenced by the historical socio-economic trends. From the unification of Italy until the early 1900’s, Lombardy’s forested land has suffered from intensive deforestation linked to demographic growth and an increasing request of timber[26, 28]. A turnaround in land management arrived when hydro-geological instability increased in deforested mountains areas. The forestry policy put forward by the Italian government selected coniferous monocultures in order to meet, together with the urge to mitigate the hydro-geological instability, the high demand of building timber [29]. More precisely, species of the genus *Pinus* have been planted in the Mediterranean region [30], while the spruce (*Picea abies*) has been widely planted in the temperate region. A decisive intervention on the reforestation of the Italian mountains has been carried out by the fascist regime whose ideology attributed to the transformation of the territory the symbol of a “national renewal and rebirth”[31]. Surveys of the 1930’s report that at the time 36 percent of the Lombardian mountain territories was occupied by forested land[28].

As a result of these historical events, the natural vegetation of the Lombardian Prealps is confined to limited areas compared to their potential range, whereas the coniferous plantations are still greatly found outside their natural range. Their presence interacts both with abiotic and biotic factors, determining an alteration of the soils properties and of biodiversity[23]. The coexistence of the two habitats - the natural vegetation and the spruce plantation - in the same region, finally provides the conditions to conduct our study and answer our research questions.

1. What is the spruce plantation’s impact on plant diversity?
2. Does the spruce plantation impact environmental factors like luminosity and ground surface temperature?
3. Do these environmental factors affect plant species occurrence and abundance?

Knowing that vegetation affects and responds to soil properties[23], we hypothesize that the spruce plantation causes a reduction of plant diversity. We also expect that the evergreen canopy of the spruce trees blocks solar radiation reducing the ground surface temperature and the undergrowth luminosity of the habitat. Finally, we imagine that plant richness positively responds to the two environmental factors, so if these decrease in the spruce plantation so does plant diversity. Indeed, luminosity and temperature regulate plant growth and phenology and are presumably responsible, together with the soil properties, for plant diversity variability between the spruce plantation and the natural habitats.

## 3 Materials and Methods

### 3.1 Geographical Setting

To pursue the analysis presented in this study case, we have collected all our original data in two different sites of the Comasche Prealps: Mount Bisbino, Cernobbio (Como) and Alpe del Vicer’e, Albavilla (Como).

The choice of the sites was, first of all, based on logistic reasons. From the University of Milan, both sites can be reached by car in less than two hours and both locations are provided with paved roads which allow almost immediate access to the habitats of interest for our study. Secondly, the two sites were chosen because they match the scientific prerequisites for our study.

The Alpe del Vicer’e - also known as Mount Bollettone[32] - and Mount Bisbino are two of the main peaks of the Como Prealps, a subsection of the Lugano Prealps that extends from the Province of Como in Italy to the Canton Ticino in Switzerland[33]. What interests us of the geographical area is the calcareous composition of the rock substrate[32, 34], which is one of the site requirements we are seeking for our study. As a matter of fact, the composition of the rock substrate affects the soil pH which is an important parameter that influences plant growth. Carbonate rocks increase the soil pH giving the soil basic properties. Consequently, the effect of the substrate of our geographical setting on soil pH is the opposite of the one of a spruce plantation’s whose litter of needles and resin cause soil acidification[23]. Finally, the two selected sites satisfy our other main requirement, which is the presence of spruce plantation patches adjacent or close to other natural or semi-natural habitats. As explained in the previous chapter, the presence of spruce plantations creates spots of acid soils in a geographical region where the rock substrate and the natural habitats would usually cause soils to have from basic to slightly acid properties (pH range = 7.7 - 6.6) [35].

Today’s presence of the spruce plantations in Mount Bisbino and the Alpe del Vicer’e is the result of the Italian fascist regime’s environmental policy that during the twentieth century has shaped the national mountain landscape[31]. These two mountain areas were once dedicated to seasonal alpine pasture activities. Their terrain used to be characterized by the presence of meadows and grazing lands that are today reduced in number and size for they have been replaced by forests and artefacts[32, 36]. During the First World War, Mount Bisbino became the southernmost stronghold of the “Frontiera Nord nel Lario Intelvese”. A military road, two artillery stakeouts and a long trench - known nowadays as “Linea Cadorna” - were built between 1916 and 1917. The landscape was ultimately modified by the natural progression of the trees on the abandoned grasslands and the artificial reforestation that began in the first half of the twentieth century[36, 37, 38]. Meanwhile, the structure of the Alpe del Vicer’e landscape was greatly changed in the 1930s when a campsite of the Opera Nazionale Balilla - an Italian Fascist youth organization[39] - was built and later converted in a big Alpine holiday village at the expenses of the grazing activities[32]. During the Second World War, the village was destroyed by an Anglo-American bombing for it had become the residence of the Accademia Navale, Accademia Aeronautica and Italian SS. In the years that followed, the war-torn site’s landscape was ultimately modified by more recent land management and administrative activities[32].

### 3.2 Data Collection

Our field activity began in March and ended by the beginning of July. The plan for our data collection initially consisted in placing a total of 15 study plots per site. In every site, the plots were meant to be organised as follows: 5 in the spruce plantation habitat and 5 in each of the other two treatments, the mixed forest and the grassland-pasture. However, we did not select any grassland-pasture plot in the Alpe del Viceŕe site because here semi-natural grassland-pastures occur at much higher elevation than forests, while the non-forested areas are lawns heavily used by human activities other than mowing and grazing (i.e., adventure park, picnic area etc.). In total, the final number of study plots from which we have collected our data is 15 in the Monte Bisbino site and 10 in the Alpe del Vicer’e site. The dimensions that we have chosen for the plots are 9 m^2^. With the objective of replicating our surveys on a monthly basis, we have obtained a total of 4 temporal replicates per study plot.

All the study plots were selected on our first two field trips, during which we have collected all their related stationary data: geographical coordinates, elevation, aspect and exposure.

To obtain the records for our plant diversity analysis, we have conducted a vegetation survey by considering the plant species occurrence and abundance in every plot. For each temporal replicate we have created an excel table where the plant species are arranged in columns and the plots in lines, the entries represent the species occurrence and are filled with 1 in case of presence and 0 in case of absence. We were not always able to identify the plant species on the field, so we have taken pictures and samples of some plants which we have identified later during data processing. Replicating our surveys in different months has been useful to identify some of the species whose recognition required to wait for the plant to bloom.

I have organized the information regarding the plant functional forms in a similar way to how we have arranged the plant species data: I have created occurrence tables where the plant functional forms are arranged in columns and the plots in lines.

Finally, all luminosity and ground surface temperature records were collected on field where we had at our disposal a light meter (Brand: Chauvin Arnoux, Model: CA 1110, Accuracy: 0.01lx, Measurement Range: 0.01lx to 200000lx) and an infrared thermometer(Brand: Bosch, Model: UniversalTemp, Optics: 12:1, Measurement Range: *−*30*^◦^C* to +500*^◦^C*). The measurement units we have used for the two parameters are respectively lux (luminous flux per unit area [40]) and degrees Celsius.

### 3.3 Data Processing

After we had completed all our field surveys, we have summarized all the collected data in a single data frame. For this operation we have compiled an excel table in which all the single plot records are arranged in lines, while in columns there are all the stationary data of our plots and all the variables we need for our analysis: the habitat type, the temporal replicate, the number of plant species, the number of plant functional forms, luminosity and temperature.

To obtain the records for the number of plant species, we have used the preliminary excel tables containing the plant species occurrence data. As a support tool for plant identification we have started a project on the website iNaturalist [41]. Here we have uploaded the plant pictures that we took during field work and obtained some suggestions for our species’ identifications. I have also used the Italian website Acta Plantarum [42] and the Swiss website Info Flora [43] to compare more images to our plant samples and obtain information about the species. Both sites have been useful to verify if the identified species were likely to be observed in the geographical area where we have performed our surveys. I have used this information to confirm some of the species identifications. Once we completed the plant species occurrence tables, we imported them on the software Rstudio. This software has a function which allows to perform a sum over the rows. As the objects in the rows of the plant occurrence data frame only take two values - 0 to indicate absence and 1 to indicate presence - we have used the sum function to calculate the total number of plant species in each plot. The records that we obtained through this sum operation are the number of plant species that we have stored in our final data frame.

After we had processed our survey’s samples and identified the plant species, I was able to arrange them in their functional forms classes. For this operation I have referred to the website Acta Plantarum, which relies on the functional forms classification elaborated by Christen Raunkiær [44]. Once the species were classified, I have completed the functional forms’ occurrence table that I had elaborated for the data collection. I then proceed with performing the Rstudio sum functions through which I have obtained the number of plant functional forms records that I have copied in the final data frame.

All luminosity and ground surface temperature records did not need any processing, therefore we have inserted them directly in the final data frame.

Once we had completed our data processing and obtained a final data frame that summarized all our variables, we were finally able to start elaborating the statistical analysis.

### 3.4 Statistical Analysis

The first question that we want to answer through our analysis is how plant diversity changes from the spruce plantation to the other habitat types - the mixed forest and the grassland-pasture.

To test the effects of the different habitats on plant diversity, we have used linear regression models. In these models, plant diversity is the response variable and the type of habitat is the explanatory variable.

### A (simple, one independent variable) linear regression model is

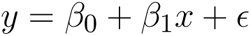

where:

- y is the response variable (e.g., plant diversity);
- x is the explanatory variable (e.g., habitat);
- *β*_1_ is the effect of x on y (i.e., the slope of the line);
- *β*_0_ is the intercept (i.e., ground or conditional mean);
- *ɛ* is the error.

The goal of fitting a regression model is to estimate the model parameters *β*_0_ and *β*_1_ given the data x and y. [45]

Plant diversity (i.e., number of plant species) has a Poisson probability distribution and our independent variable, which is the habitat type, is a categorical variable. As a consequence, we have decided to analyze plant diversity using a Generalized Linear Model (GLM) that allows us to overcome the issues of non-Normal probability distributions and of non linear independent variables.

It is possible to fit a more complex model which analyzes the interaction between explanatory variables. This model can tell us if the effect of one explanatory variable (*x*_1_) changes depending on the effect of a second explanatory variable (*x*_2_).

A linear regression model with statistical interactions is

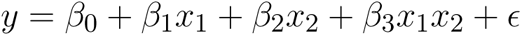

Given x and y this model allows us to estimate the *β*_0_, *β*_1_, *β*_2_ (the effect of the second explanatory variable *x*_2_) and *β*_3_ (the effect of the interaction of the two explanatory variables)[45].

We have only fitted a GLM with statistical interactions to measure the impact of the combined effects of the type of habitat (*x*_1_) and the temporal replicate (*x*_2_) on plant diversity expressed in number of species (y). When analyzing plant diversity in terms of number functional forms we have only measured the impact of the type of habitat by fitting a simple GLM. In both analysis the intercept of the model has been provided by the spruce plantation habitat.

We have fitted two other GLMs during our analysis: a single independent variable GLM to measure the effects of the type of habitat on luminosity; a GLM with statistical interactions to measure the effects of the type of habitat and the temporal replicates on the ground surface temperature. Once more, the intercept of the regression models has been provided by the spruce plantation’s habitat type.

We have performed further analysis on the two parameters to verify if luminosity and temperature have a role on determining the plant diversity trends in the different habitats. To answer this question, we have analyzed the covariance of the number of plant species and the two parameters one at a time.

Covariance is expressed as

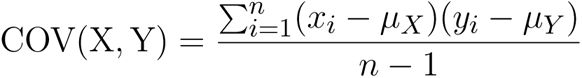

where:

- Y is the first variable (e.g., plant diversity);
- X is the second variable (e.g., luminosity, ground surface temperature);
- *µ_Y_* is the mean value of the variable Y (e.g., the mean number of species);
- *µ_X_* is the mean value of the variable X (e.g., the mean luminosity, the mean ground surface temperature).

Covariance measures the direction of a relationship between two variables. If covariance is positive we expect that if one variable increases the other variable increases too. A negative covariance indicates that as one variable increases the other one decreases. A null covariance indicates that the two variables are independent.

To finalize the analysis of the relationship between the number of species and luminosity or ground surface temperature, I have performed Pearson’s correlation test. The Pearson correlation coefficient (r) measures the strength of a linear association between two variables. A Pearson product-moment correlation attempts to draw a line of best fit through the data of two variables, and the correlation coefficient indicates how well the data points fit this line of best fit. The correlation coefficient only assumes values between +1 and *−*1: positive values indicate a positive relationship, while negative values a negative relationship; if the values are very close to +1 or *−*1, the data points almost lie on the line of best fit, on the contrary, a correlation coefficient not significantly different from 0 indicates that there is no correlation between variables.

## 4 Results

### 4.1 Plant Diversity in Number of Plant Species

We have compared three types of habitats by analyzing a total of 100 plot samples. In total we have sampled 141 plant species, 5 morphospecies within the *Poaceae, Asteraceae* and *Liliaceae* families and 1 morphospecies of fern. Our analysis reveal that land management has a significant effect on plant diversity on which the spruce plantation has a negative impact. Indeed we have observed that the number of plant species increases by 68 percent in the mixed forest compared to the spruce plantation (*β*_1_ = 0.41 *±* 0.17, P-value = 0.014). The number of plant species ultimately grows by 133 percent in the grassland-pasture habitat (*β*_1_ = 0.72 *±* 0.18, P-value *<* 0.001). This upward trend of plant diversity from the spruce plantation to the mixed forest and the grassland-pasture habitats can be well visualized from the alignment of the boxplots in figure 2.

**Figure 1:**
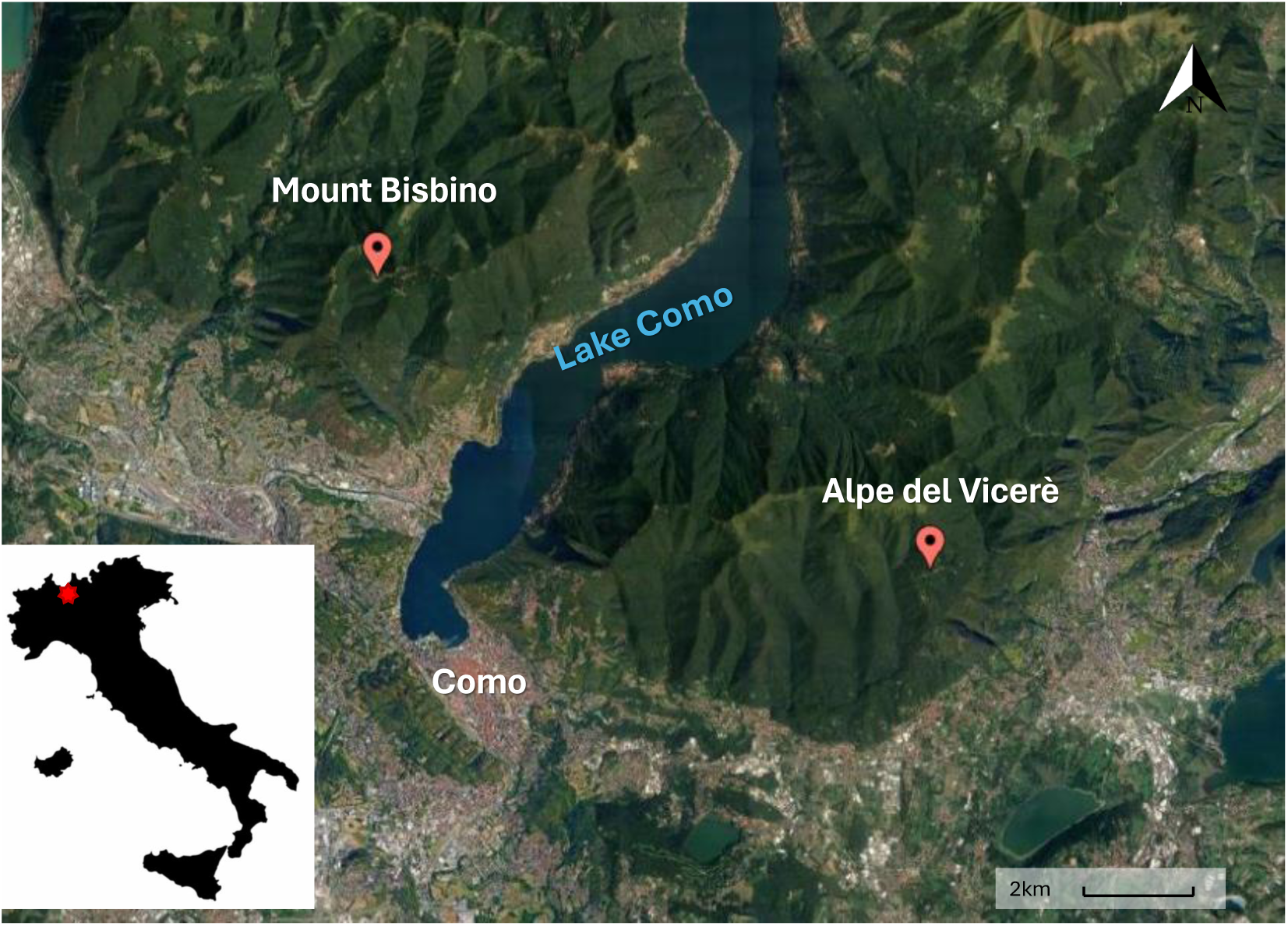
Satellite image displaying the sites’ locations.

**Figure 2:**
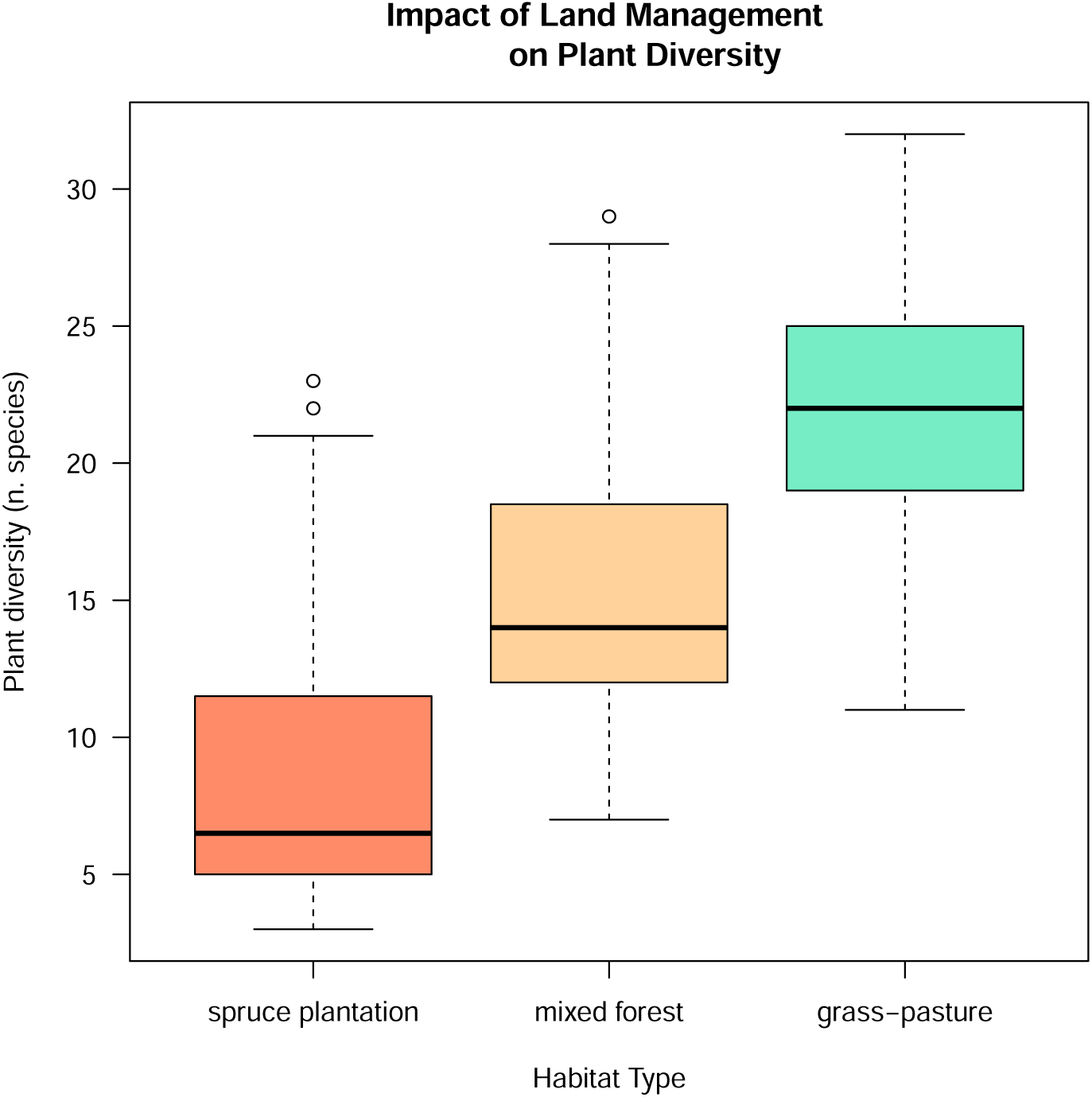
Boxplots representing the distribution of plant diversity in the three habitats: the spruce plantation, the mixed forest and the grassland-pasture (x axis). Plant diversity is here expressed in number of species (y axis).

Plant diversity did not significantly change through time nor across habitats with time (Table 1). The obtained results confirm that the number of plant species does not increase as the months of the year pass by.

**Table 1:** Number of plant species GLM summary.

|  | Coefficient | Std. Error | P-value |
| --- | --- | --- | --- |
| spruce plantation | 2.08 | 0.13 | < 0.001 |
| mixed forest | 0.41 | 0.17 | 0.014 |
| grass-pasture | 0.72 | 0.18 | < 0.001 |
| replicate | 0.06 | 0.05 | 0.173 |
| mixed forest:replicate | 0.04 | 0.06 | 0.460 |
| grass-pasture:replicate | 0.05 | 0.06 | 0.446 |

**Table 2:** Number of plant functional forms GLM summary.

|  | Coefficient | Std. Error | P-value |
| --- | --- | --- | --- |
| spruce plantation | 1.49 | 0.08 | $< 0.001$ |
| mixed forest | 0.40 | 0.10 | $< 0.001$ |
| grass-pasture | 0.30 | 0.12 | 0.013 |

**Table 3:** Luminosity GLM summary.

|  | Coefficient | Std. Error | P-value |
| --- | --- | --- | --- |
| spruce plantation | 2.01 | 0.06 | < 0.001 |
| mixed forest | 0.05 | 0.08 | 0.548 |
| grass-pasture | 0.37 | 0.09 | < 0.001 |

### 4.2 Plant Diversity in Plant Functional Forms Richness

The spruce plantation has a negative impact on the diversity of plant functional forms too. As we can observe in Figure 3, we have recorded the highest diversity values in the mixed forest. In this habitat, diversity increases by 49 percent (*β*_1_ = 0.40 *±* 0.10, P-value *<* 0.001). From the spruce plantation to the grassland-pasture the diversity of plant functional forms increases by 31 percent (*β*_1_ = 0.30 *±* 0.12, P-value = 0.013).

**Figure 3:**
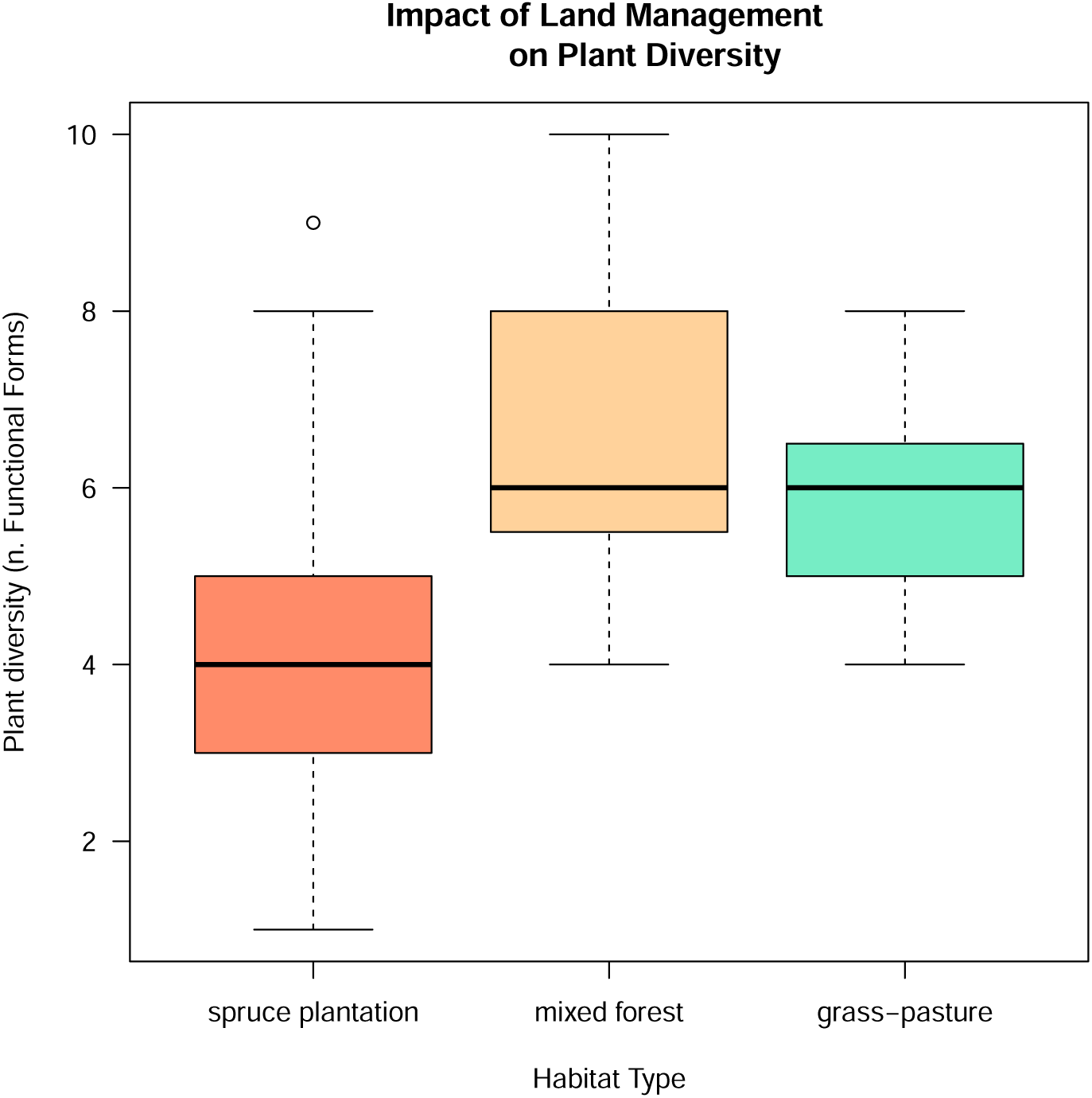
Boxplots representing the distribution of plant diversity in the three habitats: the spruce plantation, the mixed forest and the grassland-pasture (x axis). Plant diversity is here expressed in number of plant functional forms (y axis).

### 4.3 Luminosity and Temperature

The amount of luminosity that irradiates our plots is impacted by land management, we can see in figure 4 how the distribution of the collected luminosity values differs in the habitats. We have not found a significant difference between the two forest habitats of our study case, the mixed forest treatment does not significantly impact luminosity (*β*_1_ = 0.05 *±* 0.08, P-value = 0.548). However light intensity increases by 32 percent in the grassland-pasture habitat (*β*_1_ = 0.37 *±* 0.09, P-value *<* 0.001).

**Figure 4:**
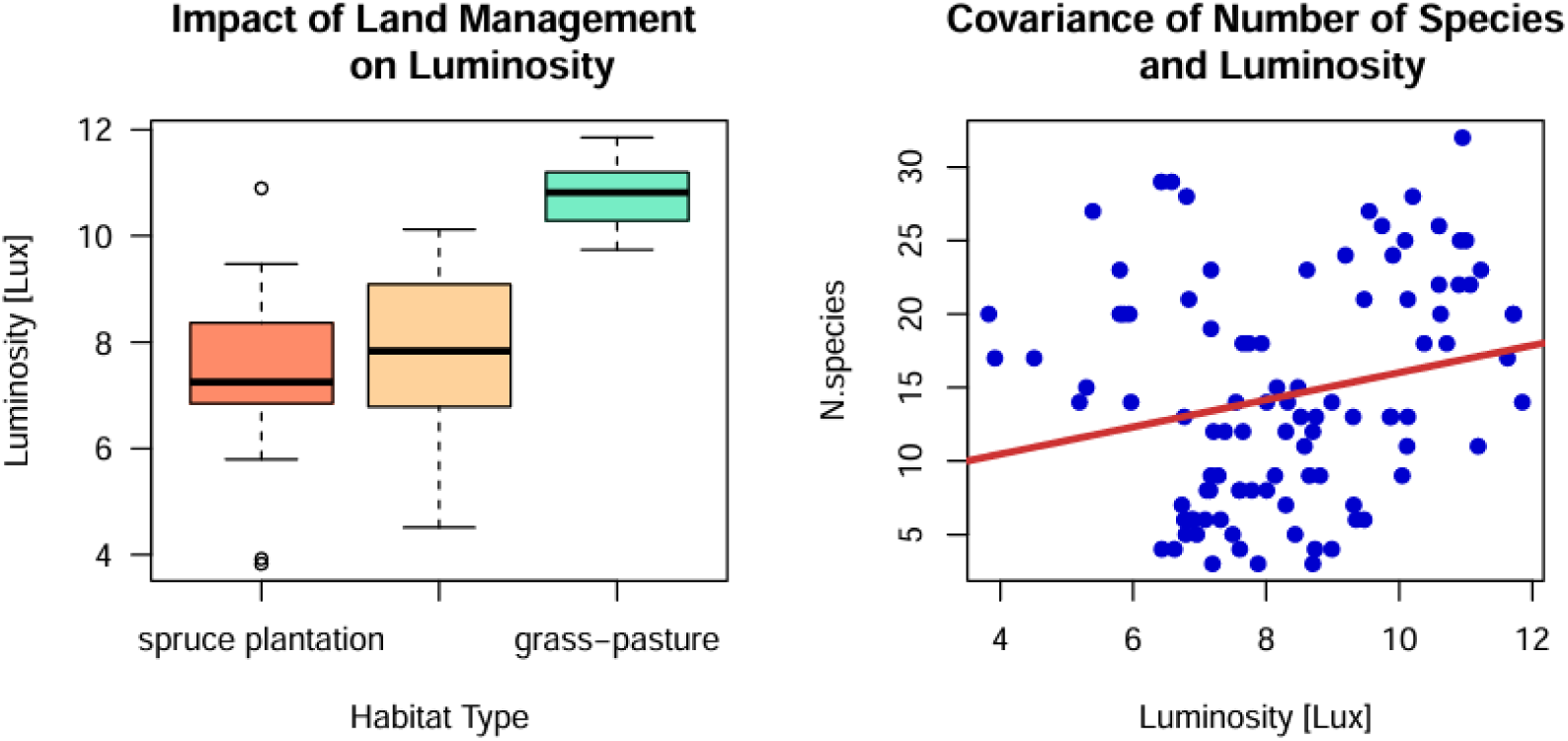
Left - Boxplots representing the distribution of luminosity in the three habitats: the spruce plantation, the mixed forest and the grassland-pasture. Right - Scatterplot displaying covariance between luminosity (x axis) and the number of species (y axis): every blue dot represents a record, regression line is represented in red.

**Figure 5:**
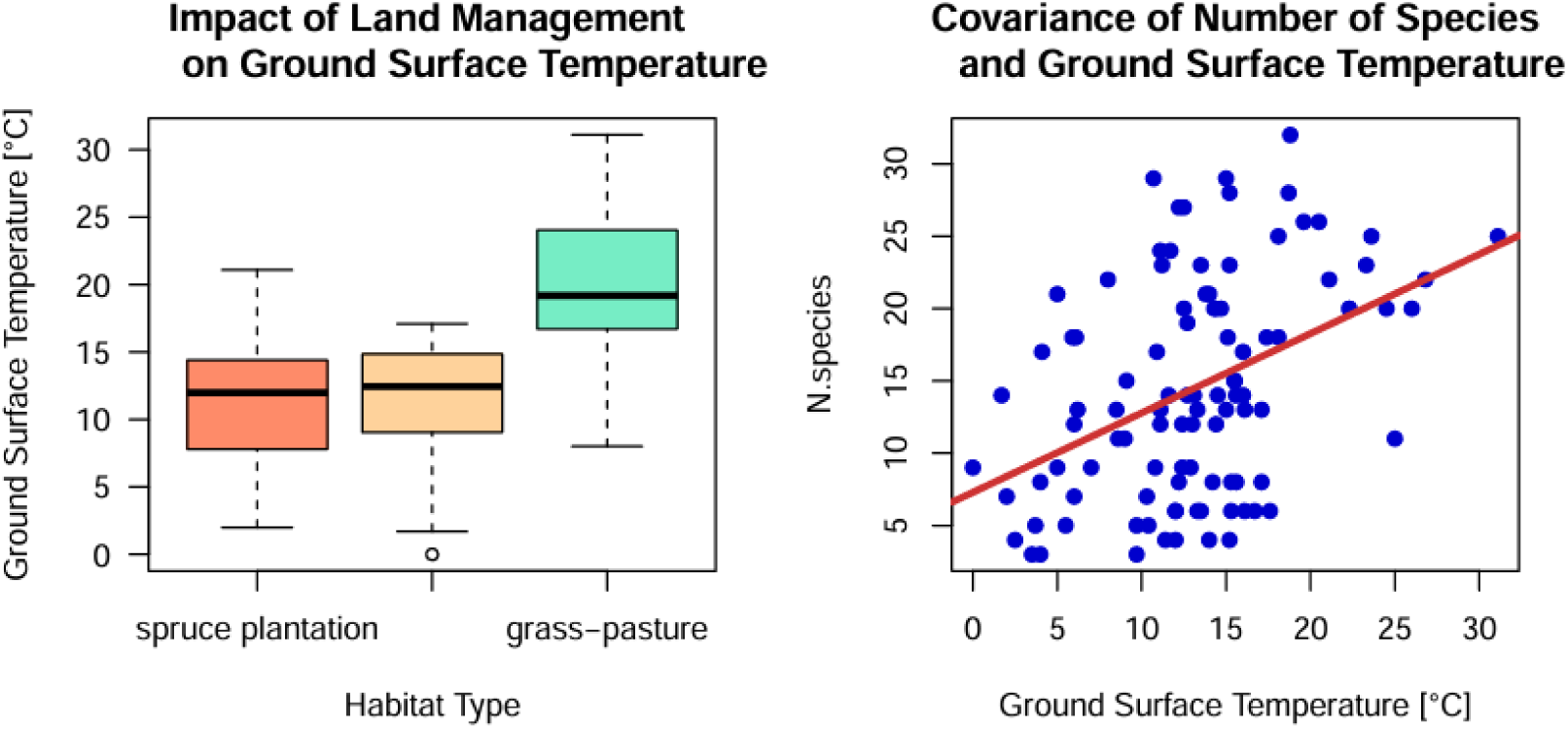
Left - Boxplots representing the distribution of ground surface temperature in the three habitats: the spruce plantation, the mixed forest and the grassland-pasture. Right - Scatterplot displaying covariance between ground surface temperature (x axis) and the number of plant species (y axis): every blue dot represents a record, regression line is represented in red.

We have observed that a positive covariance exists between luminosity and the number of plant species (cov = 3.06). The relationship between the two variables is displayed in figure 4. I have performed Pearson’s correlation test and obtained a P-value equal to 0.022, this confirms the linear correlation of our records (r = 0.26).

The impact of land management on the ground surface temperature of the habitats does not differ too much from what we have observed for luminosity - a positive relationship exists between luminosity and ground surface temperature (cov = 4.58, r = 0.43, correlation test P-value = *<* 0.001). Ground surface temperatures do not significantly differ between the spruce plantation and the mixed forest (Table 4), however they do increase by 80 percent in the grassland-pasture habitat (*β*_1_ = 1.26 *±* 0.19, P-value *<* 0.001).

**Table 4:** Ground surface temperature GLM summary.

|  | Coefficient | Std. Error | P-value |
| --- | --- | --- | --- |
| spruce plantation | 1.47 | 0.14 | $< 0.001$ |
| mixed forest | 0.22 | 0.19 | 0.241 |
| grass-pasture | 1.26 | 0.19 | $< 0.001$ |
| replicate | 0.35 | 0.04 | $< 0.001$ |
| mixed forest:replicate | -0.07 | 0.06 | 0.266 |
| grass-pasture:replicate | -0.24 | 0.06 | $< 0.001$ |

Ground surface temperature varies in time, indeed the temporal replicate variable has a significant effect on the parameter (*β*_2_ = 0.35 *±* 0.04, P-value *<* 0.001). Only the interaction of the temporal replicate with the grassland-pasture habitat has a significant effect on ground surface temperature (*β*_3_ = *−*0.24 *±* 0.06, P-value *<* 0.001) the negative coefficient reveals that as time passes the impact of the habitat type is less strong (Figure 6).

**Figure 6:**
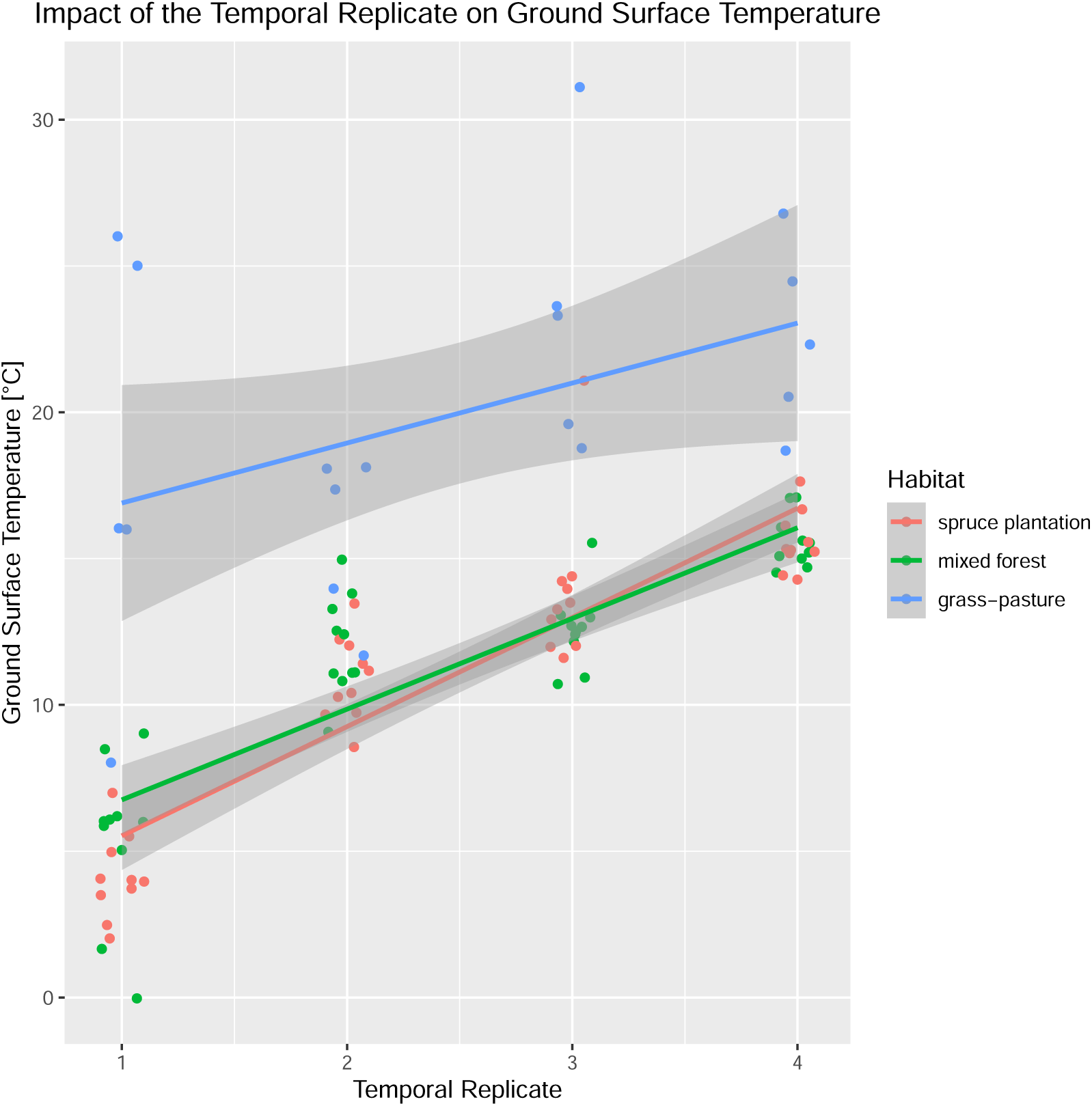
Scatterplot displaying how geound surface temperature (y-axis) varies through time, expressed in temporal replicates (x-axis), in the different habitats. Every dot represents a record, each colour a type of habitat: the spruce plantation in red, the mixed forest in green and the grassland-pasture in blue.

A positive covariance exists between ground surface temperature and the number of species too (cov = 18.5). The linear correlation of our records is confirmed by the Pearson’s correlation test (*r* = 0.42 and P-value *<* 0.001).

## 5 Discussion

### 5.1 Main Findings

We have examined the effects of land management, namely reforestation, on vegetation and understood how significant are the responses of plant diversity to the different land-use treatments. What we learn from our research is that spruce plantation causes a significant decrease of plant diversity.

With the highest mean of plant species (*µ* = 21.85), the grassland-pasture is the habitat that favours plant diversity the most, indeed grasslands are normally richer in plant species than forested lands. After spruce plantations (*µ* = 9.375) have been set up at the place of grassland-pastures, their plant diversity has decreased by 57 percent. The mixed forest - which is the most similar habitat to the natural potential vegetation of our study area - places itself in the middle of the ranking for number of species (*µ* = 15.775). Nevertheless, the fact that the number of species decreases by 41 percent from the mixed forest to the spruce plantation is a significant result.

In addition to the results obtained from the number of species, the analysis on the number of plant functional forms appoint the mixed forest as the most favourable treatment to plant diversity (*µ* = 6.6). As we could have expected, being a habitat that is subject to constant disturbance - grazing animals and mowing - the grassland-pasture has a lower mean of plant functional forms than the mixed forest (*µ* = 5.95). But the most relevant impact on diversity is once more the spruce plantation’s (*µ* = 4.425). The plantation has caused, in fact, a 21 percent decrease of plant functional forms compared to the grassland-pasture, which is the habitat that once used to cover the land of our sites. The passage from the potential vegetation of the sites, the mixed forest, to the spruce plantation also reveals a significant 33 percent decrease of plant functional forms.

From our results we learn that the only habitat which significantly impacts luminosity and ground surface temperature is the grassland-pasture, while the mixed forest and the spruce plantation have the same effect on the two parameters. To the grassland-pasture belong the highest records of luminosity and ground surface temperature and this is also the habitat with the highest number of plant species. Our results confirm that a positive relationship exists between the number of plant species and luminosity and also between the number of plant species and ground surface temperature.

Of all the response variables that we have considered, only the ground surface temperature has revealed a significant response to the influence of the temporal replicate. Our results show that ground surface temperatures increase from spring to the beginning of summer in all the habitats, however they increase differently depending on the type of habitat.

### 5.2 Implications

In the light of our findings, the spruce plantation is not a positive presence in the Como Prealps. Biodiversity is essential for supporting ecosystem functioning and its loss causes destabilization and decay[13]. It is therefore likely that the spruce plantations that we have studied are destined to collapse. Together with the reduction of plant species other symptoms are already visible, for instance, the absence of young spruce specimen or seedlings and the presence of diseased trees are signs that the spruce plantation is no longer a resilient habitat.

In parallel, land management ought to preserve grasslands, for they represent the habitat that favours plant species richness the most. In addition, grasslands could also contribute to climate change mitigation. Indeed, they have a higher albedo range (0.16 - 0.26) than forest habitats (0.15 - 0.20 for deciduous forests, 0.05 - 0.15 for coniferous forests)[46]. Albedo is the percentage of the total solar incoming radiation that is reflected by a surface[47]. The presence of grasslands among forests would help increase the total albedo rate contributing to climate change mitigation while preserving plant diversity.

The absence of age variability between the spruce specimen combined with the reduction of plant functional forms in the spruce plantation indicate that land management has perturbed the forest’s structure. The vertical structure of the forest affects energy transfer and biomass production[48], indeed structural heterogeneity allows a greater exploitation of light and nutrients[49, 50]. Secondly, structural diversity creates a series of micro-habitats that will on their turn favour biodiversity[49]. A greater presence of herbaceous plants and shrubs, together with adult trees, confers the mixed forest a more complex vertical structure that is absent or poorly represented in the spruce plantation. Thus, the mixed forest is a richer, more stable and better functioning habitat.

Plant species variety and functional forms diversity imply temporal variability in the phenology of the plants. Phenology is an important feature that interests us in this study because it is responsible for the co-existence of certain species and can therefore explain the plant richness differences between the three habitats. As a fact, we know that a plant’s phenology is influenced by environmental factors like temperature and luminosity and that different species adapt their life cycle depending on how the parameters vary in the habitat they live in. For this reason, there are some geophyte plants characterized by an early flowering time in spring that we have found in the grassland-pasture and the mixed forest - dominated by deciduous trees - but not in the spruce plantation - dominated by evergreen trees [51]. Species like *Narcissus poeticus*, *Scilla bifolia* and *Crocus versicolor* (figure 7) are examples of geophytes that have an early flowering season to anticipate the leafing of the other plants of the habitat that causes a reduction of luminosity on the ground. We have not found these species in the spruce plantation habitat where the canopy blocks luminosity all year long.

**Figure 7:**
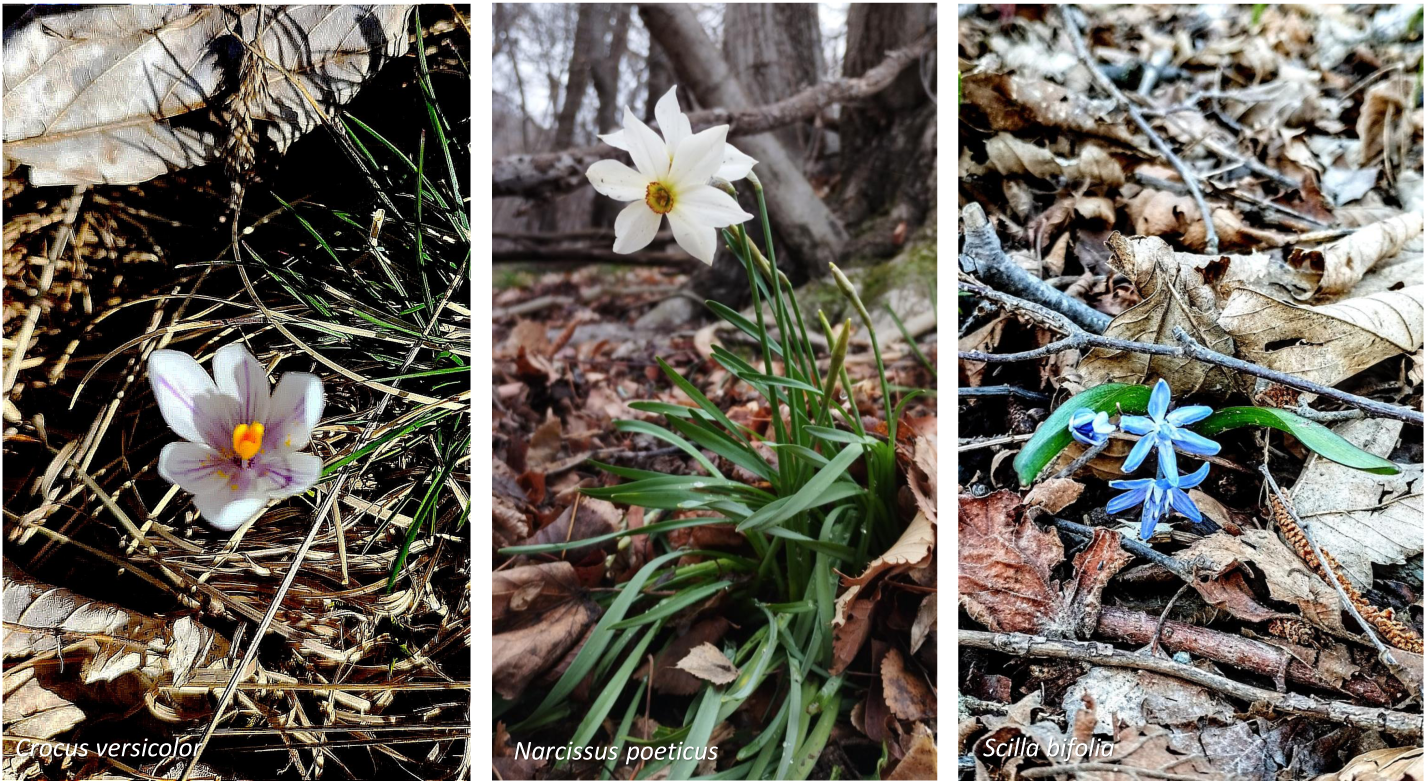
Pictures of three geophyte plants taken in the grassland-pasture and mixed forest plots.

Unfortunately, the analysis of our records does not reveal that the interaction of the temporal replicates with the mixed forest habitat has a significant impact on luminosity and ground surface temperature. Nevertheless, the two parameters highly depend on the season and the daily weather, so it is possible that a higher survey frequency and a year long data collection will provide different results.

What we can confirm through our results is that luminosity and ground surface temperature are influenced by the land-use treatment and highly increase in the grassland-pasture, the habitat with the higher number of plant species. Their implication in determining plant diversity is intuitive, for instance we can imagine that the absence of a tree canopy in the grassland-pasture reduces the competition for sunlight allowing more plants to grow. However, the positive impact of the parameters on the number of species has also been confirmed by the correlation tests that we have performed.

Once more, our results on luminosity and ground surface temperature fail to explain why plant species increase from the spruce plantation to the mixed forest. It is likely that other parameters play a role on determining biodiversity, among these, the soil pH would be an interesting variable to measure. Further studies on the environmental factors that influence plant growth and phenology - e.g., luminosity, temperature, soil pH etc. - will provide a clearer overview of the causes of biodiversity loss in a monoculture. Among these, future studies could investigate how animal diversity responds to spruce plantation and examine effective restoration ideas.

### 5.3 Conclusions

This research provides us important evidence that an incorrect land management has relevant and long-lasting consequences on plant diversity. What we learn is that in order to preserve plant diversity and ecosystem functioning, monocultures are best to be avoided. At their place, we suggest that future reforestation projects consider planting a higher diversity of exclusively native tree species with the aim of reproducing the natural potential vegetation of the habitat. If the semi-natural vegetation favours biodiversity more than the natural vegetation - like the grassland-pasture does in our case of study - it would be a good idea to include both habitats in the land management programmes. For instance, if we think of our case in the Como Prealps, the reforestation of the abandoned grasslands should also imply the creation and maintenance of some grass patches among the matrix of mixed forest.

Another concern that arises from our results is finding a solution to convert the already existing monocultures into mixed-species stands. With this objective, we have already started an experiment in the Alpe del Vicer’e site. In a spruce plantation spot of the site we have planted some tree seedlings of native species like *Fraxinus excelsior*, *Castanea sativa* and *Abies alba* together with other native shrubs. The growth of these plants is monitored together with their impact on the plantation’s biodiversity.

In current times, the realization of climate change consequences is bringing a lot of attention on the urge to reduce carbon emissions. Monocultures can appear as the quickest solution to absorb carbon from the atmosphere, for instance, coniferous plantations have been promoted by forest managers for their relatively simple management among other economical reasons [15]. But, as our research demonstrates, on the long term monocultures have negative consequences on biodiversity and ecosystem functioning that we must not ignore. As a matter of facts, there is increasing evidence that species-rich plantations stock much more carbon than monocultures. Notably, biodiversity loss and climate change are two intertwined processes, one can cause the other and vice-versa [25]. It is very unfortunate that a unified international Convention on both climate change mitigation and biodiversity protection does yet not exist[12]. Indeed, if climate change mitigation programmes are not readjusted in order to protect biodiversity, their success will only be temporary and precarious, while their damage might be irreversible.

Returning to our concern on the impact of reforestation, our research demonstrates what can be the long term consequences of incorrect land management. From our research we learn that it would be more advisable to encompass biodiversity protection measures in future reforestation-based projects aimed at climate change mitigation. For instance, these measures should include creation and maintenance of a higher variability of habitats - e.g., grass patches among the matrix of forests, which increase plant diversity and surface albedo - and investment in mixed-species plantations that will guarantee the development of healthy, long lasting and highly efficient ecosystems.

## Acknowledgments

We are thankful to ERSAF – Ente Regionale per i Servizi all’Agricoltura e alle Foreste – of Region Lombardy and to the Municipality of Albavilla for allowing the implementation of this research.

## A Appendix

**Table 5:**
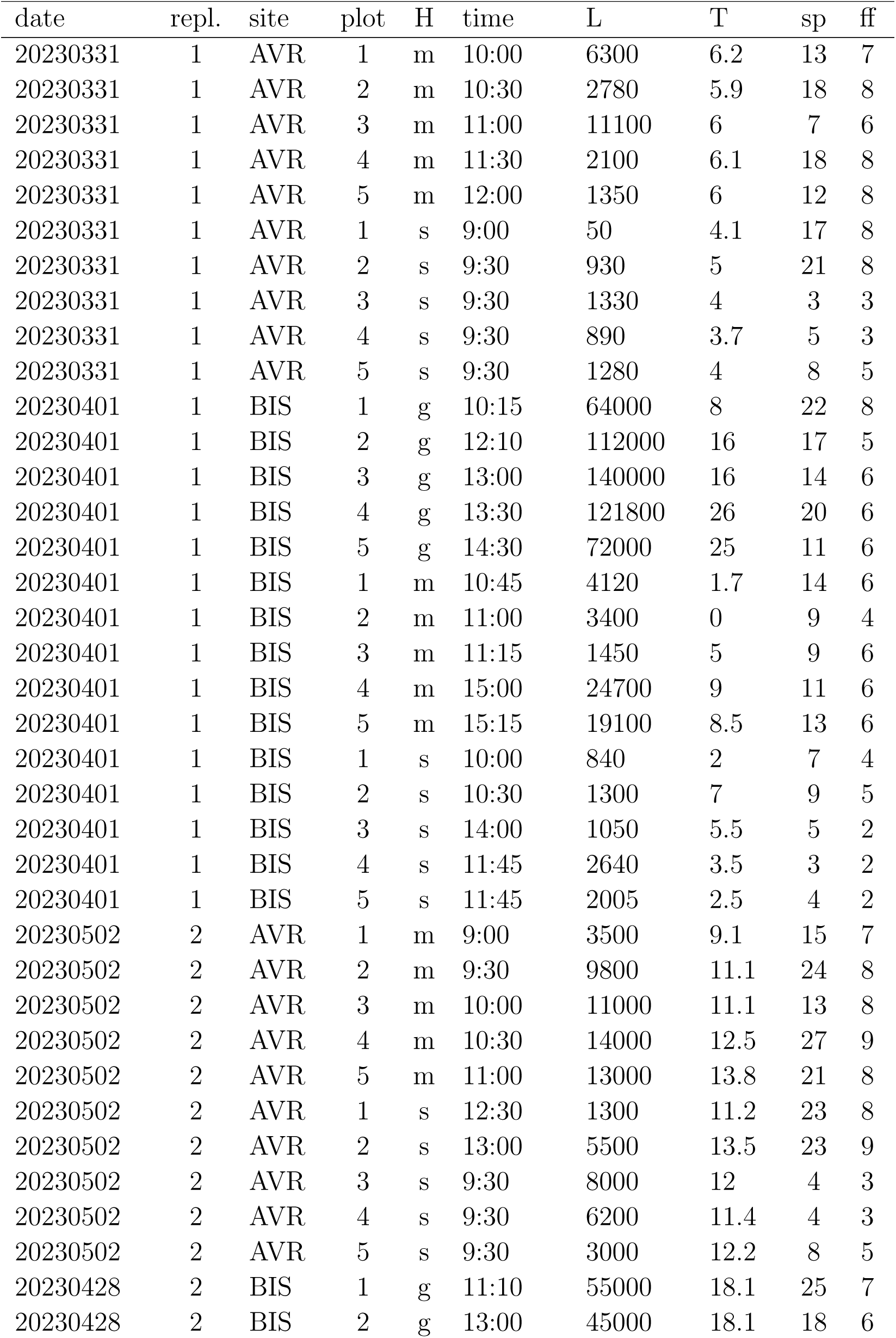
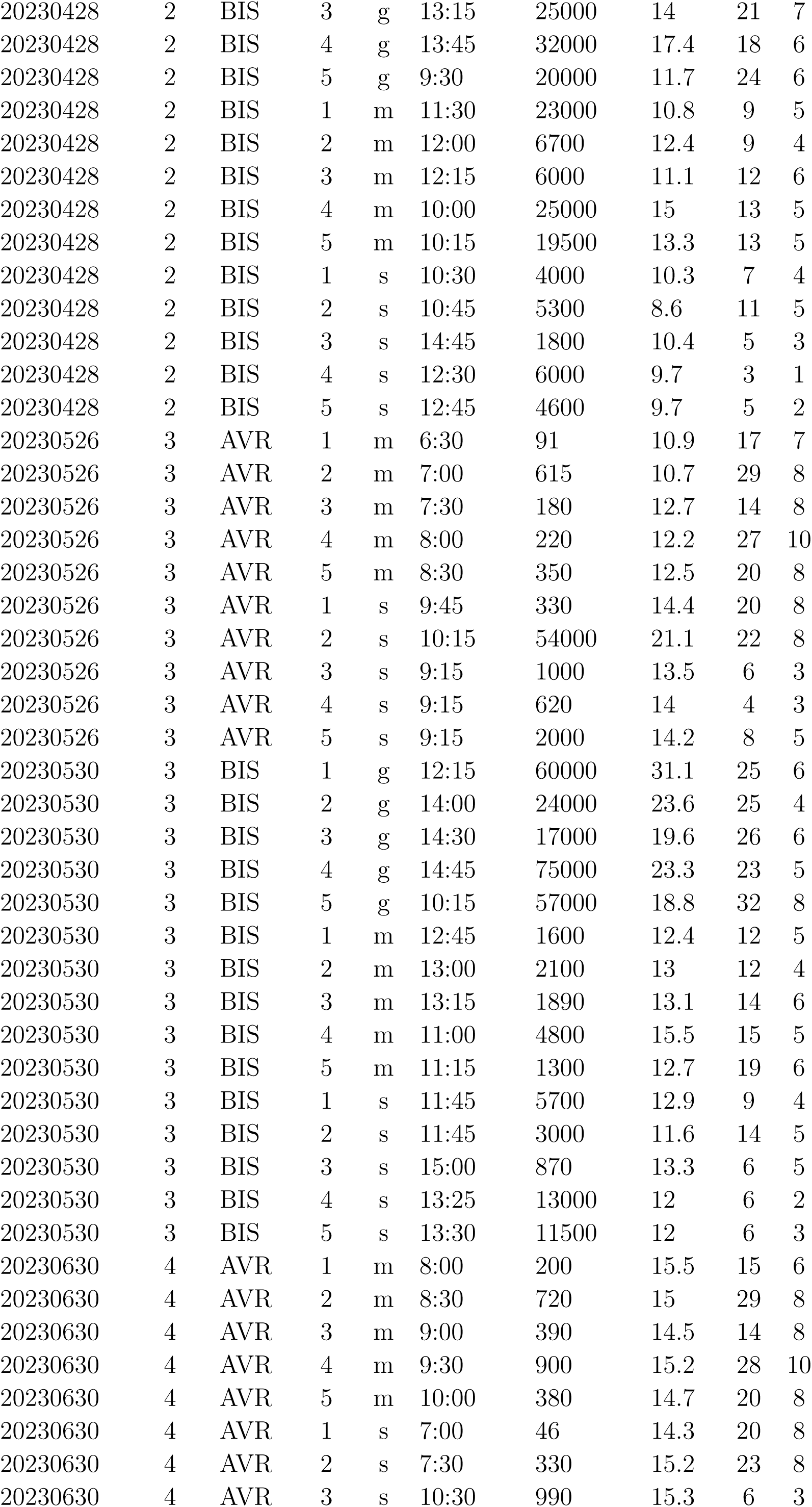
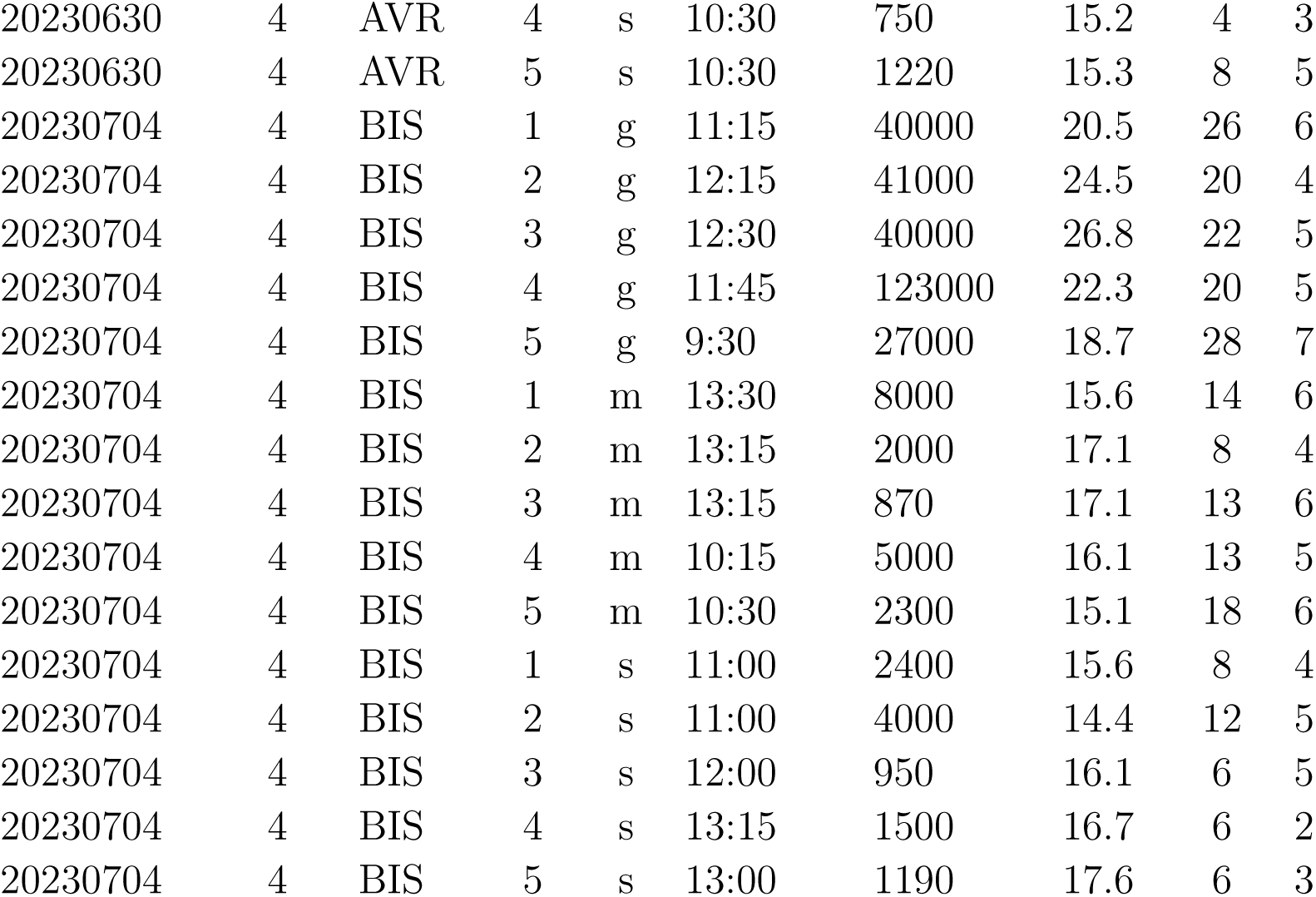
Sampling Data Frame. H: habitat. L: luminosity [Lux]. T: ground surface temperature [°C]. sp: number of plant species. ff: number of plant functional forms. Site: AVR = Alpe del Viceŕe, BIS = Monte Bisbino. Habitat: g = grass-pasture, m = mixed forest, s = spruce plantation.

| date | repl. | site | plot | H | time | L | T | sp | ff |
| --- | --- | --- | --- | --- | --- | --- | --- | --- | --- |
| 20230331 | 1 | AVR | 1 | m | 10:00 | 6300 | 6.2 | 13 | 7 |
| 20230331 | 1 | AVR | 2 | m | 10:30 | 2780 | 5.9 | 18 | 8 |
| 20230331 | 1 | AVR | 3 | m | 11:00 | 11100 | 6 | 7 | 6 |
| 20230331 | 1 | AVR | 4 | m | 11:30 | 2100 | 6.1 | 18 | 8 |
| 20230331 | 1 | AVR | 5 | m | 12:00 | 1350 | 6 | 12 | 8 |
| 20230331 | 1 | AVR | 1 | s | 9:00 | 50 | 4.1 | 17 | 8 |
| 20230331 | 1 | AVR | 2 | s | 9:30 | 930 | 5 | 21 | 8 |
| 20230331 | 1 | AVR | 3 | s | 9:30 | 1330 | 4 | 3 | 3 |
| 20230331 | 1 | AVR | 4 | s | 9:30 | 890 | 3.7 | 5 | 3 |
| 20230331 | 1 | AVR | 5 | s | 9:30 | 1280 | 4 | 8 | 5 |
| 20230401 | 1 | BIS | 1 | g | 10:15 | 64000 | 8 | 22 | 8 |
| 20230401 | 1 | BIS | 2 | g | 12:10 | 112000 | 16 | 17 | 5 |
| 20230401 | 1 | BIS | 3 | g | 13:00 | 140000 | 16 | 14 | 6 |
| 20230401 | 1 | BIS | 4 | g | 13:30 | 121800 | 26 | 20 | 6 |
| 20230401 | 1 | BIS | 5 | g | 14:30 | 72000 | 25 | 11 | 6 |
| 20230401 | 1 | BIS | 1 | m | 10:45 | 4120 | 1.7 | 14 | 6 |
| 20230401 | 1 | BIS | 2 | m | 11:00 | 3400 | 0 | 9 | 4 |
| 20230401 | 1 | BIS | 3 | m | 11:15 | 1450 | 5 | 9 | 6 |
| 20230401 | 1 | BIS | 4 | m | 15:00 | 24700 | 9 | 11 | 6 |
| 20230401 | 1 | BIS | 5 | m | 15:15 | 19100 | 8.5 | 13 | 6 |
| 20230401 | 1 | BIS | 1 | s | 10:00 | 840 | 2 | 7 | 4 |
| 20230401 | 1 | BIS | 2 | s | 10:30 | 1300 | 7 | 9 | 5 |
| 20230401 | 1 | BIS | 3 | s | 14:00 | 1050 | 5.5 | 5 | 2 |
| 20230401 | 1 | BIS | 4 | s | 11:45 | 2640 | 3.5 | 3 | 2 |
| 20230401 | 1 | BIS | 5 | s | 11:45 | 2005 | 2.5 | 4 | 2 |
| 20230502 | 2 | AVR | 1 | m | 9:00 | 3500 | 9.1 | 15 | 7 |
| 20230502 | 2 | AVR | 2 | m | 9:30 | 9800 | 11.1 | 24 | 8 |
| 20230502 | 2 | AVR | 3 | m | 10:00 | 11000 | 11.1 | 13 | 8 |
| 20230502 | 2 | AVR | 4 | m | 10:30 | 14000 | 12.5 | 27 | 9 |
| 20230502 | 2 | AVR | 5 | m | 11:00 | 13000 | 13.8 | 21 | 8 |
| 20230502 | 2 | AVR | 1 | s | 12:30 | 1300 | 11.2 | 23 | 8 |
| 20230502 | 2 | AVR | 2 | s | 13:00 | 5500 | 13.5 | 23 | 9 |
| 20230502 | 2 | AVR | 3 | s | 9:30 | 8000 | 12 | 4 | 3 |
| 20230502 | 2 | AVR | 4 | s | 9:30 | 6200 | 11.4 | 4 | 3 |
| 20230502 | 2 | AVR | 5 | s | 9:30 | 3000 | 12.2 | 8 | 5 |
| 20230428 | 2 | BIS | 1 | g | 11:10 | 55000 | 18.1 | 25 | 7 |
| 20230428 | 2 | BIS | 2 | g | 13:00 | 45000 | 18.1 | 18 | 6 |
| 20230428 | 2 | BIS | 3 | g | 13:15 | 25000 | 14 | 21 | 7 |
| 20230428 | 2 | BIS | 4 | g | 13:45 | 32000 | 17.4 | 18 | 6 |
| 20230428 | 2 | BIS | 5 | g | 9:30 | 20000 | 11.7 | 24 | 6 |
| 20230428 | 2 | BIS | 1 | m | 11:30 | 23000 | 10.8 | 9 | 5 |
| 20230428 | 2 | BIS | 2 | m | 12:00 | 6700 | 12.4 | 9 | 4 |
| 20230428 | 2 | BIS | 3 | m | 12:15 | 6000 | 11.1 | 12 | 6 |
| 20230428 | 2 | BIS | 4 | m | 10:00 | 25000 | 15 | 13 | 5 |
| 20230428 | 2 | BIS | 5 | m | 10:15 | 19500 | 13.3 | 13 | 5 |
| 20230428 | 2 | BIS | 1 | s | 10:30 | 4000 | 10.3 | 7 | 4 |
| 20230428 | 2 | BIS | 2 | s | 10:45 | 5300 | 8.6 | 11 | 5 |
| 20230428 | 2 | BIS | 3 | s | 14:45 | 1800 | 10.4 | 5 | 3 |
| 20230428 | 2 | BIS | 4 | s | 12:30 | 6000 | 9.7 | 3 | 1 |
| 20230428 | 2 | BIS | 5 | s | 12:45 | 4600 | 9.7 | 5 | 2 |
| 20230526 | 3 | AVR | 1 | m | 6:30 | 91 | 10.9 | 17 | 7 |
| 20230526 | 3 | AVR | 2 | m | 7:00 | 615 | 10.7 | 29 | 8 |
| 20230526 | 3 | AVR | 3 | m | 7:30 | 180 | 12.7 | 14 | 8 |
| 20230526 | 3 | AVR | 4 | m | 8:00 | 220 | 12.2 | 27 | 10 |
| 20230526 | 3 | AVR | 5 | m | 8:30 | 350 | 12.5 | 20 | 8 |
| 20230526 | 3 | AVR | 1 | s | 9:45 | 330 | 14.4 | 20 | 8 |
| 20230526 | 3 | AVR | 2 | s | 10:15 | 54000 | 21.1 | 22 | 8 |
| 20230526 | 3 | AVR | 3 | s | 9:15 | 1000 | 13.5 | 6 | 3 |
| 20230526 | 3 | AVR | 4 | s | 9:15 | 620 | 14 | 4 | 3 |
| 20230526 | 3 | AVR | 5 | s | 9:15 | 2000 | 14.2 | 8 | 5 |
| 20230530 | 3 | BIS | 1 | g | 12:15 | 60000 | 31.1 | 25 | 6 |
| 20230530 | 3 | BIS | 2 | g | 14:00 | 24000 | 23.6 | 25 | 4 |
| 20230530 | 3 | BIS | 3 | g | 14:30 | 17000 | 19.6 | 26 | 6 |
| 20230530 | 3 | BIS | 4 | g | 14:45 | 75000 | 23.3 | 23 | 5 |
| 20230530 | 3 | BIS | 5 | g | 10:15 | 57000 | 18.8 | 32 | 8 |
| 20230530 | 3 | BIS | 1 | m | 12:45 | 1600 | 12.4 | 12 | 5 |
| 20230530 | 3 | BIS | 2 | m | 13:00 | 2100 | 13 | 12 | 4 |
| 20230530 | 3 | BIS | 3 | m | 13:15 | 1890 | 13.1 | 14 | 6 |
| 20230530 | 3 | BIS | 4 | m | 11:00 | 4800 | 15.5 | 15 | 5 |
| 20230530 | 3 | BIS | 5 | m | 11:15 | 1300 | 12.7 | 19 | 6 |
| 20230530 | 3 | BIS | 1 | s | 11:45 | 5700 | 12.9 | 9 | 4 |
| 20230530 | 3 | BIS | 2 | s | 11:45 | 3000 | 11.6 | 14 | 5 |
| 20230530 | 3 | BIS | 3 | s | 15:00 | 870 | 13.3 | 6 | 5 |
| 20230530 | 3 | BIS | 4 | s | 13:25 | 13000 | 12 | 6 | 2 |
| 20230530 | 3 | BIS | 5 | s | 13:30 | 11500 | 12 | 6 | 3 |
| 20230630 | 4 | AVR | 1 | m | 8:00 | 200 | 15.5 | 15 | 6 |
| 20230630 | 4 | AVR | 2 | m | 8:30 | 720 | 15 | 29 | 8 |
| 20230630 | 4 | AVR | 3 | m | 9:00 | 390 | 14.5 | 14 | 8 |
| 20230630 | 4 | AVR | 4 | m | 9:30 | 900 | 15.2 | 28 | 10 |
| 20230630 | 4 | AVR | 5 | m | 10:00 | 380 | 14.7 | 20 | 8 |
| 20230630 | 4 | AVR | 1 | s | 7:00 | 46 | 14.3 | 20 | 8 |
| 20230630 | 4 | AVR | 2 | s | 7:30 | 330 | 15.2 | 23 | 8 |
| 20230630 | 4 | AVR | 3 | s | 10:30 | 990 | 15.3 | 6 | 3 |
| 20230630 | 4 | AVR | 4 | s | 10:30 | 750 | 15.2 | 4 | 3 |
| 20230630 | 4 | AVR | 5 | s | 10:30 | 1220 | 15.3 | 8 | 5 |
| 20230704 | 4 | BIS | 1 | g | 11:15 | 40000 | 20.5 | 26 | 6 |
| 20230704 | 4 | BIS | 2 | g | 12:15 | 41000 | 24.5 | 20 | 4 |
| 20230704 | 4 | BIS | 3 | g | 12:30 | 40000 | 26.8 | 22 | 5 |
| 20230704 | 4 | BIS | 4 | g | 11:45 | 123000 | 22.3 | 20 | 5 |
| 20230704 | 4 | BIS | 5 | g | 9:30 | 27000 | 18.7 | 28 | 7 |
| 20230704 | 4 | BIS | 1 | m | 13:30 | 8000 | 15.6 | 14 | 6 |
| 20230704 | 4 | BIS | 2 | m | 13:15 | 2000 | 17.1 | 8 | 4 |
| 20230704 | 4 | BIS | 3 | m | 13:15 | 870 | 17.1 | 13 | 6 |
| 20230704 | 4 | BIS | 4 | m | 10:15 | 5000 | 16.1 | 13 | 5 |
| 20230704 | 4 | BIS | 5 | m | 10:30 | 2300 | 15.1 | 18 | 6 |
| 20230704 | 4 | BIS | 1 | s | 11:00 | 2400 | 15.6 | 8 | 4 |
| 20230704 | 4 | BIS | 2 | s | 11:00 | 4000 | 14.4 | 12 | 5 |
| 20230704 | 4 | BIS | 3 | s | 12:00 | 950 | 16.1 | 6 | 5 |
| 20230704 | 4 | BIS | 4 | s | 13:15 | 1500 | 16.7 | 6 | 2 |
| 20230704 | 4 | BIS | 5 | s | 13:00 | 1190 | 17.6 | 6 | 3 |

**Table 6:** Collected data summary.

| **Variable** | **Habitat** | **min** | **MAX** | **mean($\mu$)** |
| --- | --- | --- | --- | --- |
| Number of plant species | spruce plantation | 3 | 23 | 9.375 |
|  | mixed forest | 7 | 29 | 15.775 |
|  | grass-pasture | 11 | 32 | 21.85 |
| Number of plant functional forms | spruce plantation | 1 | 9 | 4.425 |
|  | mixed forest | 4 | 10 | 6.6 |
|  | grass-pasture | 4 | 8 | 5.95 |
| Luminosity [Lux] | spruce plantation | 46 | 54000 | 4085.29 |
|  | mixed forest | 91 | 25000 | 6045.15 |
|  | grass-pasture | 17000 | 140000 | 59540 |
| Ground Surface Temp. [°C] | spruce plantation | 2 | 21.1 | 11.125 |
|  | mixed forest | 0 | 17.1 | 11.405 |
|  | grass-pasture | 8 | 31.1 | 19.975 |

**Table 7:** Generalized Linear Models summary.

| **Model** | **Explanatory variable** | $\beta$ | $\epsilon$ | **p-value** |
| --- | --- | --- | --- | --- |
| Number of plant species | spruce plantation | 2.08 | 0.13 | < 0.001 |
|  | mixed forest | 0.41 | 0.17 | 0.014 |
|  | grass-pasture | 0.72 | 0.18 | < 0.001 |
|  | replicate | 0.06 | 0.05 | 0.173 |
|  | mixed forest:replicate | 0.04 | 0.06 | 0.460 |
|  | grass-pasture:replicate | 0.05 | 0.06 | 0.446 |
| Number of plant functional forms | spruce plantation | 1.49 | 0.08 | < 0.001 |
|  | mixed forest | 0.40 | 0.10 | < 0.001 |
|  | grass-pasture | 0.30 | 0.12 | 0.013 |
| Luminosity | spruce plantation | 2.01 | 0.06 | < 0.001 |
|  | mixed forest | 0.05 | 0.08 | 0.548 |
|  | grass-pasture | 0.37 | 0.09 | < 0.001 |
| Ground surface temperature | spruce plantation | 1.47 | 0.14 | < 0.001 |
|  | mixed forest | 0.22 | 0.19 | 0.241 |
|  | grass-pasture | 1.26 | 0.19 | < 0.001 |
|  | replicate | 0.35 | 0.04 | < 0.001 |
|  | mixed forest:replicate | -0.07 | 0.06 | 0.266 |
|  | grass-pasture:replicate | -0.24 | 0.06 | < 0.001 |

**Table 8:**
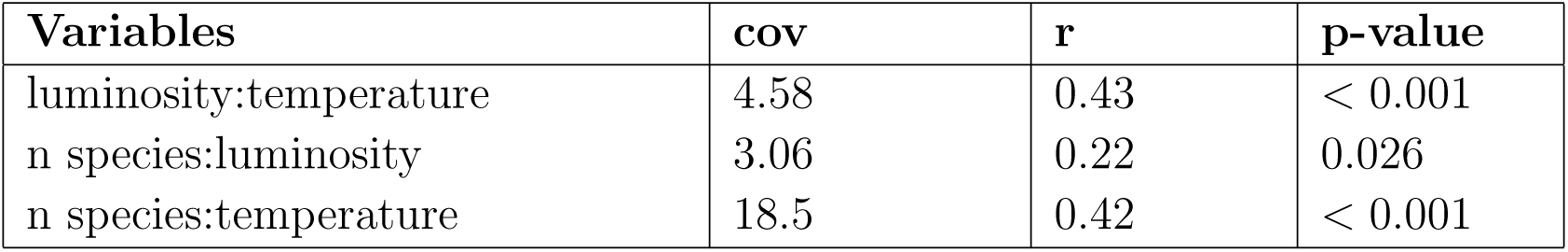
Covariance values and Pearson’s correlation coefficients and P-values.

| Variables | cov | r | p-value |
| --- | --- | --- | --- |
| luminosity:temperature | 4.58 | 0.43 | < 0.001 |
| n species:luminosity | 3.06 | 0.22 | 0.026 |
| n species:temperature | 18.5 | 0.42 | < 0.001 |

**Table 9:** Sampled Species List with functional form class and number of plots containing the species per each habitat. S = spruce plantation, M = mixed forest, G = grass-pasture

| Plant Species | Functional Form | S | M | G |
| --- | --- | --- | --- | --- |
| *Abies alba* | P scap | 3 | 0 | 0 |
| *Acer campestre* | P scap | 1 | 0 | 0 |
| *Acer pseudoplatanus* | P scap | 2 | 5 | 1 |
| *Achillea millefolium* | H scap | 0 | 0 | 5 |
| *Adoxa moschatellina* | G rhiz | 0 | 3 | 0 |
| *Agrostis sp* | H caesp | 0 | 0 | 4 |
| *Ajuga sp* | NA | 0 | 0 | 3 |
| *Alchemilla sp* | H ros | 0 | 0 | 2 |
| *Alliaria petiolata* | H scap | 2 | 5 | 0 |
| *Anemonoides nemorosa* | G rhiz | 2 | 6 | 2 |
| *Angelica sp* | H scap | 0 | 1 | 0 |
| *Anthoxanthum odoratum* | H caesp | 0 | 0 | 2 |
| *Aquilegia sp* | H scap | 0 | 1 | 0 |
| *Aristolochia pallida* | G bulb | 0 | 1 | 0 |
| *Arrhenatherum elatius* | H caesp | 0 | 0 | 5 |
| *Arum sp* | G rhiz | 0 | 2 | 0 |
| *Asperula taurina* | G rhiz | 0 | 3 | 0 |
| *Astrantia minor* | H scap | 0 | 0 | 1 |
| *Betonica officinalis* | H scap | 0 | 0 | 5 |
| *Brachipodium sylvaticum* | H caesp | 1 | 0 | 2 |
| *Briza media* | H caesp | 0 | 0 | 4 |
| *Bromus erectus* | H caesp | 0 | 0 | 4 |
| *Cardamine sp* | G rhiz | 0 | 1 | 0 |
| *Cardamine bulbifera* | G rhiz | 0 | 1 | 0 |
| *Cardamine heptaphylla* | G rhiz | 2 | 1 | 0 |
| *Carduus sp* | NA | 0 | 0 | 3 |
| *Carex sp* | H caesp | 1 | 0 | 3 |
| *Castanea sativa* | P scap | 0 | 5 | 0 |
| *Centaurea jacea* | H scap | 0 | 0 | 1 |
| *Clematis sp* | NA | 0 | 0 | 1 |
| *Clematis vitalba* | P lian | 3 | 4 | 0 |
| *Clinopodium menthifolium* | H scap | 0 | 1 | 0 |
| *Convolvulus sp* | G rhiz | 1 | 1 | 0 |
| *Cornus sp* | P caesp | 0 | 2 | 0 |
| *Corylus avellana* | P caesp | 0 | 5 | 0 |
| *Crocus sp* | G bulb | 0 | 1 | 0 |
| *Crocus versicolor* | G bulb | 0 | 0 | 2 |
| *Cruciata glabra* | H scap | 0 | 0 | 5 |
| *Cyclamen purpurascens* | G bulb | 0 | 2 | 0 |
| *Cynosurus cristatus* | H caesp | 0 | 0 | 1 |
| *Cytisus insubricus* | P caesp | 0 | 1 | 0 |
| *Dactylis glomerata* | H caesp | 1 | 2 | 5 |
| *Daphne laureola* | P caesp | 2 | 0 | 0 |
| *Digitalis lutea* | H scap | 0 | 1 | 0 |
| *Erigeron sp* | NA | 0 | 1 | 0 |
| *Erythronium dens-canis* | G bulb | 2 | 2 | 0 |
| *Euphorbia dulcis* | G rhiz | 1 | 0 | 0 |
| *Euphrasia sp* | T scap | 0 | 0 | 3 |
| *Fagus sylvatica* | P scap | 3 | 5 | 0 |
| *Festuca sp* | H caesp | 0 | 0 | 3 |
| *Fraxinus excelsior* | P scap | 6 | 6 | 0 |
| *Galium sp* | H scap | 1 | 1 | 0 |
| *Galium lucidum* | H scap | 0 | 0 | 5 |
| *Galium odoratum* | G rhiz | 1 | 2 | 1 |
| *Geranium nodosum* | G rhiz | 2 | 3 | 0 |
| *Geranium robertianum* | T scap | 3 | 1 | 0 |
| *Geum sp* | H ros | 0 | 2 | 0 |
| *Hedera helix* | P lian | 5 | 4 | 0 |
| *Helleborus viridis* | G rhiz | 2 | 1 | 0 |
| *Hepatica nobilis* | G rhiz | 2 | 0 | 0 |
| *Hieracium sp* | H ros | 2 | 1 | 0 |
| *Hieracium pilosella* | H ros | 0 | 0 | 5 |
| *Hypericum sp* | NA | 0 | 1 | 3 |
| *Hippocrepis comosa* | H caesp | 0 | 0 | 5 |
| *Ilex aquifolium* | P caesp | 2 | 1 | 0 |
| *Knautia drymeja* | H scap | 0 | 0 | 4 |
| *Laburnum alpinum* | P scap | 3 | 0 | 0 |
| *Larix decidua* | P scap | 1 | 0 | 0 |
| *Lathyrus sp* | NA | 0 | 1 | 0 |
| *Lathyrus pratensis* | H scap | 0 | 0 | 5 |
| *Leontodon sp* | H ros | 0 | 0 | 1 |
| *leucanthemum sp* | H scap | 0 | 0 | 3 |
| *Lilium sp* | G bulb | 0 | 1 | 0 |
| *Lotus sp* | NA | 0 | 0 | 4 |
| *Luzula sp* | H caesp | 1 | 3 | 0 |
| *Luzula campestris* | H caesp | 0 | 0 | 4 |
| *Luzula multiflora* | H caesp | 0 | 0 | 1 |
| *Luzula nivea* | H caesp | 4 | 5 | 0 |
| *Melica sp* | H caesp | 0 | 1 | 0 |
| *Melica nutans* | H caesp | 0 | 1 | 0 |
| *Milium effusum* | H caesp | 2 | 3 | 0 |
| *Molinia caerulea* | H caesp | 0 | 0 | 4 |
| *Mycelis muralis* | H scap | 1 | 2 | 0 |
| *Narcissus poeticus* | G bulb | 0 | 4 | 1 |
| *Ornithogalum pyrenaicum* | G bulb | 0 | 1 | 0 |
| *Ostrya carpinifolia* | P scap | 0 | 1 | 0 |
| *Oxalis acetosella* | G rhiz | 1 | 5 | 0 |
| *Phleum rhaeticum* | H scap | 0 | 0 | 1 |
| *Phyteuma scheuchzeri* | H scap | 0 | 0 | 3 |
| *Picea abies* | P scap | 9 | 0 | 0 |
| *Pimpinella sp* | H scap | 0 | 1 | 1 |
| *Plantago altissima* | H ros | 0 | 0 | 2 |
| *Poa sp* | H scap | 0 | 0 | 2 |
| *Poa pratensis* | H caesp | 0 | 0 | 2 |
| *Polygala comosa* | H scap | 0 | 0 | 4 |
| *Polygonatum sp* | G rhiz | 0 | 3 | 0 |
| *Polygonatum odoratum* | G rhiz | 1 | 3 | 0 |
| *Potentilla sp* | NA | 1 | 4 | 1 |
| *Potentilla erecta* | H scap | 0 | 0 | 5 |
| *Prenantes sp* | H scap | 0 | 1 | 0 |
| *Primula sp* | H ros | 1 | 1 | 1 |
| *Primula vulgaris* | H ros | 2 | 0 | 0 |
| *Prunus avium* | P scap | 2 | 1 | 0 |
| *Prunus laurocerasus* | P scap | 0 | 1 | 0 |
| *Pseudoturritis turrita* | H scap | 0 | 1 | 0 |
| *Ranunculus acris* | H scap | 0 | 0 | 5 |
| *Ranunculus platanifolius* | H scap | 0 | 4 | 0 |
| *Rhinanthus alectorolophus* | T scap | 0 | 0 | 1 |
| *Robinia pseudoacacia* | P scap | 0 | 2 | 0 |
| *Rosa sp* | P caesp | 1 | 0 | 1 |
| *Rubus sp* | NA | 2 | 3 | 1 |
| *Rubus saxatilis* | H scap | 4 | 5 | 0 |
| *Rumex acetosa* | H scap | 0 | 0 | 3 |
| *Salvia glutinosa* | H scap | 0 | 1 | 0 |
| *Salvia pratensis* | H scap | 0 | 0 | 1 |
| *Sanguisorba officinalis* | H scap | 0 | 0 | 3 |
| *Scilla bifolia* | G bulb | 1 | 4 | 0 |
| *Senecio sp* | H scap | 1 | 2 | 0 |
| *Senecio nemorensis* | H scap | 0 | 3 | 0 |
| *Sorbus aria* | P scap | 0 | 3 | 0 |
| *Sorbus aucuparia* | P scap | 1 | 1 | 0 |
| *Stellaria sp* | H scap | 0 | 1 | 0 |
| *Stellaria graminea* | H scap | 0 | 0 | 4 |
| *Stellaria nemorum* | H scap | 5 | 5 | 3 |
| *Symphytum tuberosum* | G rhiz | 0 | 1 | 0 |
| *Tanacetum sp* | H scap | 0 | 1 | 0 |
| *Taraxacum sp* | H ros | 0 | 1 | 0 |
| *Teucrium chamaedrys* | Ch suffr | 0 | 0 | 1 |
| *Thymus sp* | Ch rept | 0 | 0 | 5 |
| *Tilia cordata* | P scap | 1 | 3 | 0 |
| *Trifolium pratense* | H scap | 0 | 0 | 4 |
| *Trifolium repens* | H rept | 0 | 0 | 4 |
| *Urtica dioica* | H scap | 1 | 0 | 0 |
| *Veratrum album* | G rhiz | 0 | 4 | 0 |
| *veronica chamaedrys* | H scap | 0 | 0 | 5 |
| *Veronica persica* | T scap | 0 | 1 | 0 |
| *Vicia sepia* | H scap | 0 | 0 | 2 |
| *Vinca minor* | Ch rept | 2 | 3 | 0 |
| *Viola sp* | H scap | 2 | 4 | 0 |
| *Viola hirta* | H ros | 0 | 0 | 1 |
| *Viola reichenbachiana* | H scap | 1 | 0 | 5 |

## Notes

### Competing Interest Statement

The authors have declared no competing interest.

